# VBayesMM: Variational Bayesian neural network to prioritize important relationships of high-dimensional microbiome multiomics data

**DOI:** 10.1101/2024.11.27.625587

**Authors:** Tung Dang, Artem Lysenko, Keith A. Boroevich, Tatsuhiko Tsunoda

## Abstract

The analysis of high-dimensional microbiome multiomics datasets is crucial for understanding the complex interactions between microbial communities and host physiological states across health and disease conditions. Despite their importance, current methods such as the microbe–metabolite vectors (MMvec) approach, often fail to efficiently identify keystone species. This arises from the vast dimensionality of metagenomics data which complicates the inference of significant relationships, particularly the estimation of co-occurrence probabilities between microbes and metabolites. Here we propose the variational Bayesian microbiome multiomics (VBayesMM) approach, which enhances MMvec by incorporating a spike-and-slab prior within a Bayesian neural network. This allows VBayesMM to rapidly and precisely identify crucial microbial species, improving the accuracy of estimated co-occurrence probabilities between microbes and metabolites, while also robustly managing the uncertainty inherent in high-dimensional data. Moreover, we have implemented variational inference to address computational bottlenecks, enabling scalable analysis across extensive multiomics datasets. Our comparative evaluation of large-scale human and mouse multiomics datasets demonstrates that VBayesMM not only outperforms existing methods in accuracy but also provides a scalable solution for analyzing massive datasets. VBayesMM enhances the interpretability of the Bayesian neural network by identifying a core set of influential microbial species, thus facilitating a deeper understanding of their probabilistic relationships with the host.

## 1 Introduction

Recent studies have shown that shifts in the composition of the human gut microbiome significantly influence overall health [1, 2] and contribute to the development of various diseases such as inflammatory bowel disease [3], and cancer [4–7]. By leveraging advanced metagenomic sequencing techniques such as 16S rRNA sequencing and shotgun metagenomics, scientists can identify microbial species linked to various diseases and investigate their functional roles within microbial communities [8–10]. The integration of diverse omics technologies — metagenomics, transcriptomics, and metabolomics — further enriches our understanding of the dynamic interactions between the host and its microbiome in human health and disease. Gut metabolites are primarily produced or influenced by microbial enzymes that metabolize dietary elements and substances secreted by the host. Key microbial metabolites, such as short-chain fatty acids (SCFAs) and bile acids, serve as vital indicators of microbial fermentation and metabolism processes [11, 12]. For instance, the combination of low colonic SCFAs and high bile acids correlates with an increased risk of colon cancer among Americans consuming diets high in fats and proteins but low in complex carbohydrates [13]. Thus, the combined application of metagenomics and metabolomics presents a promising approach to decipher the complex biochemical interactions between the gut microbiota and the host [14, 15].

To explore the integration of disparate omics data, particularly the combination of microbial sequencing with mass spectrometry technologies for metabolites, researchers have developed several integrative analysis methods. However, current approaches encounter notable challenges when handling multiomics data. One primary issue arises from the differing measurement scales between sequencing and mass spectrometry, which complicates the analysis of relationships between microbiomes and metabolites due to the need for scale invariance [16, 17]. Traditional analytical methods, such as canonical correspondence analysis and sparse partial least squares (sPLS) regression [18, 19], struggle to maintain scale invariance when applied to inter-related datasets, primarily due to their underlying assumption of independent relationships. This limitation can lead to potential inaccuracies when combining microbiome and mass spectrometry data [16]. Furthermore, integrating these technologies produce high-dimensional and compositional datasets, making it challenging to discern biologically meaningful connections among the vast array of diverse biological variables. To tackle these challenges, neural network-based approaches have been developed to estimate co-occurrence probabilities between microbes and metabolites, offering a more robust solution for capturing complex, non-linear relationships in multiomics data. The microbe–metabolite vectors (MMvec) approach, implemented in QIIME 2, is one of the most widely used methods for exploring microbiome-metabolite relationships [16, 20]. MMvec employs neural networks to calculate the conditional probabilities of detecting specific metabolites given the presence of particular microbes, thus addressing the limitations of previous methods that treated microbe–metabolite relationships as independent. This approach more effectively captures the complex interdependencies in the data. Additionally, the neural network architecture of MMvec is specifically designed to handle the compositional nature of microbiome and mass spectrometry datasets. Traditional methods often struggle with inconsistencies between absolute and relative abundances across these datasets. By ensuring consistency between these two abundances, MMvec effectively reduces the false discovery rates [16]. The practical significance of the MMvec approach is well-documented, with multiple applications in human health and disease [21–23], environmental studies [24], and animal microbiome [25, 26], highlighting its utility in exploring the potential interrelations between microbiomes and metabolites across diverse fields.

However, the vast dimensionality of microbial metagenomics datasets poses significant challenges for the MMvec approach. Within this framework, all taxonomic units (microbial species) are treated with equal importance when computing co-occurrence probabilities between microbes and metabolites. This assumption, however, may not hold in practice, as a large number of taxonomic units might be irrelevant and thus may not contribute meaningfully to the identification or characterization of representative microbe-metabolite relationships. Emerging methods such as microbiome-metabolome network (MiMeNet) [27] or MIMOSA2 [28] have used neural network-based approaches similar to MMvec, further enhancing the exploration of microbe–metabolite relationships. These methods, which utilize regularization techniques like *L*_2_ regularization to address issues of sparsity [29], still face challenges in optimizing the choice of hyperparameters and in providing interpretable context for selected taxonomic units. Thus, there is still a definite need for an interpretable selection of key representative taxonomic units and probabilistically inferring which types of metabolites they are associated with. Furthermore, while the maximum a posteriori (MAP) probability estimates used in MMvec and standard approaches such as the sPLS and MiMeNet methods provide valuable point estimates for weight matrices and bias vectors, they do not sufficiently capture the uncertainty inherent in such predictive models, which can result in overconfident predictions that are ultimately inaccurate. These inaccuracies can result in valuable experimental resources being wasted on investigating ineffective or erroneous compounds [30–32]. Therefore, incorporating a reliable representation of uncertainty in predictive models is crucial to mitigate these risks. Recently, simulation approaches such as Markov chain Monte Carlo (MCMC) methods have been explored to improve uncertainty quantification in deep learning models, which could potentially enhance the reliability of their predictions [33, 34]. However, the direct applications of these approaches to microbiome multiomics studies present significant challenges due to the computational burden and the difficulty of achieving convergence in such complex high-dimensional datasets [35, 36].

In this study, we propose VBayesMM (variational Bayesian microbiome multiomics), a Bayesian neural network designed to address these limitations and enhance computational scalability for large microbiome multiomics datasets. The main contributions of this study are threefold. First, to overcome MMvec’s limitations in accurately assessing the importance of taxonomic units, we implement a spike-and-slab prior approach [37–39]. Unlike standard regularization techniques, the spike-and-slab approach improves feature selection by probabilistically distinguishing between relevant and irrelevant taxonomic units. This method enhances our ability to detect a minimal core set of taxonomic units that contribute to representative co-occurrence probabilities between microbes and metabolites. By incorporating spike-and-slab priors, we enable a probabilistic interpretation of each taxonomic unit’s relevance, thereby improving interpretability, which is further supported by also offering methods to visualize the probabilistic significance of each feature.

Second, to manage uncertainty and model complexity in neural networks, we adopt a Bayesian neural network that incorporates uncertainties in weights and bias [40, 41]. This probabilistic framework allows us to calculate the standard deviation of the posterior probabilities of weight matrices and bias vectors, effectively quantifying predictive uncertainty. VBayesMM not only enhances the robustness of the network’s predictions but also provides a reliable measure of predictive uncertainty.

Third, to address the computational inefficiencies traditionally associated with Bayesian methods when applied in high-dimensional settings, we implement a variational inference method [42, 43]. This approach transforms the challenging high-dimensional sampling tasks relied on by methods like the MCMC into a more manageable optimization problem. Variational inference has proven effective across a wide range of applications, including the analysis of large datasets related to metagenomics [44, 45], population genetics [46], and single cells [47–49].

Finally, to validate the performance of VBayesMM, we selected several high-quality published microbiome datasets profiled using 16S rRNA gene amplicon and whole genome shotgun sequencing methods, which were paired with mass spectrometry metabolite composition profiling [16, 50–52]. Specifically, we used two published 16S rRNA gene amplicon sequencing datasets: one from a study on mice with obstructive sleep apnea, containing 4,690 taxonomic units with 382 metabolite abundances [50], and another from a study examining the effects of a high-fat diet, containing 913 taxonomic units with 2,257 metabolite abundances [16]. Additionally, we used larger datasets from whole genome shotgun sequencing studies on gastric [52] and colorectal cancers [51], which include approximately 40,000 and 50,000 taxonomic units, respectively. These diverse datasets allowed us to validate the proposed method in a range of distinctive high-complexity settings and to rigorously test its computational efficiency and scalability.

## 2 Results

### 2.1 Overview of Variational Bayesian microbiome multiomics (VBayesMM) approach

We propose the development of the VBayesMM approach (Fig. 1 and Methods), building on our previously introduced variational Bayesian method [42], which was shown to be highly effective for the tasks of microbiome clustering [44] and analyses of microbiome-metabolite relationships [45]. The main contributions in this study are threefold: First, the VBayesMM approach focuses on estimating the conditional probabilities of observing specific metabolites given the presence of taxonomic units (or microbial taxa). This approach allows the probabilistic identification and prioritization of a small number of microbiome-metabolite relationships from high-dimensional datasets, allowing potential applications in the study of human microbiome-related diseases. Second, the VBayesMM framework is structured as a variational Bayesian neural network [40, 41], which captures the uncertainties inherent in microbiome and metabolite datasets derived from sequencing technologies and mass spectrometry. These uncertainties arise from a mixture of biological and technical factors. Finally, VBayesMM handles the vast dimensionality of input data, including the relative abundances of microbial species, to generate accurate predictions of metabolite abundance profiles.

**Fig. 1:**
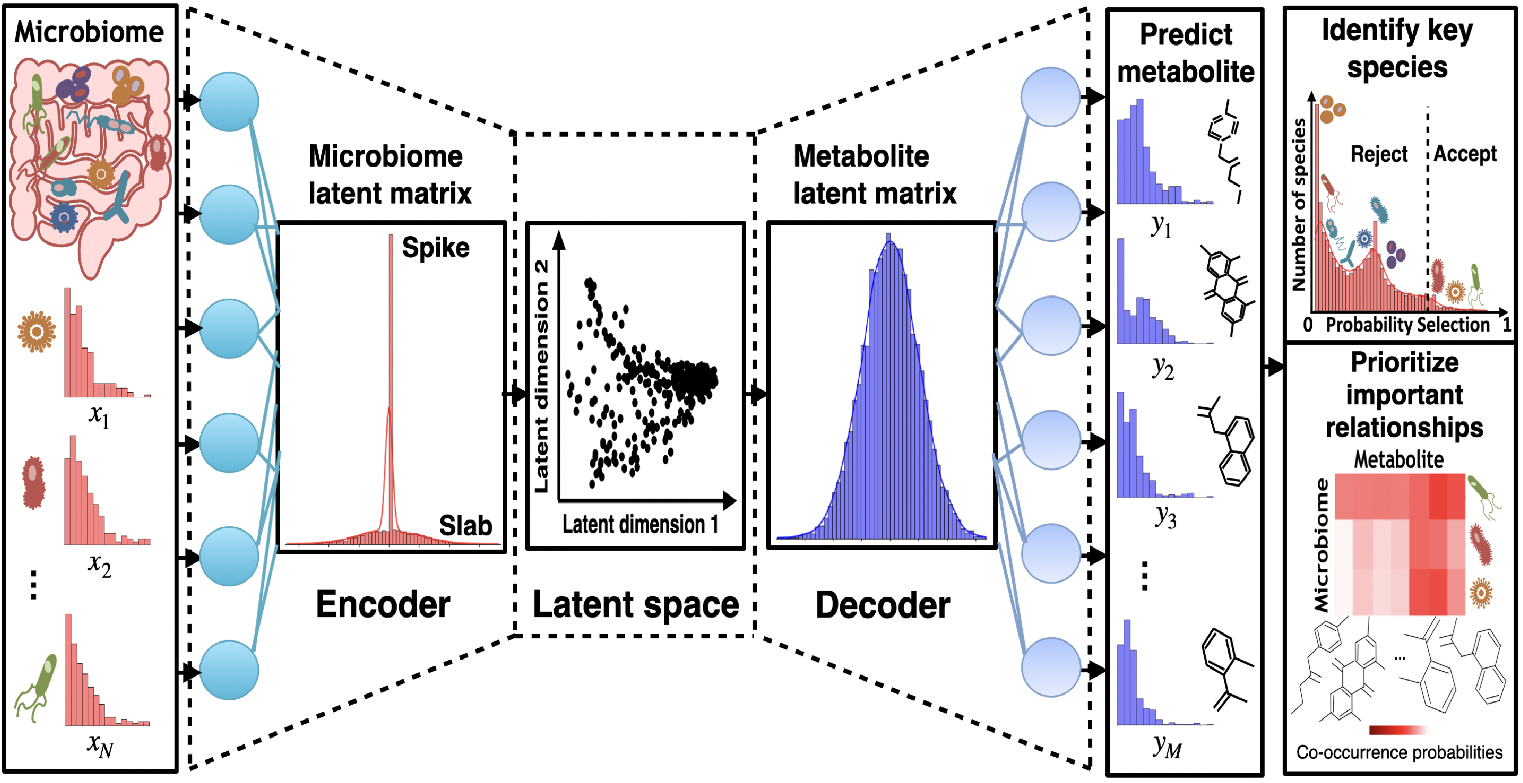
VBayesMM neural network architecture. The VBayesMM approach uses paired microbiome-metabolite data, with microbial species as input variables and metabolite abundances as target variables. VBayesMM encodes input microbial sequence counts (x) (using neural network encoder) into a lower-dimensional latent space which is utilized by the decoder to estimate conditional probabilities of observing specific metabolite abundances (y). VBayesMM implements a variational spike-and-slab distribution within the microbiome latent matrix to improve the probabilistic identification of microbial species, aiming to enhance predictive accuracy. A subset of microbial species, selected based on probabilistic thresholds, is then integrated with the metabolite latent matrix to explore microbiome-metabolite relationships. These relationships can be measured and ranked by the co-occurrence probabilities. Variational inference and stochastic optimization techniques are applied to effectively manage the computational challenges posed by the vast dimensionality of the input data.

As shown in figure 1, the encoder component of the neural network learns the multidimensional probability distribution of these inputs, projecting them into a lower-dimensional latent space based on observed microbial sequence counts. To isolate a small subset of microbial species (or taxonomic units) that enhance the accuracy and relevance of these predictions, we propose using a variational spike-and-slab distribution within the encoder neural network (applied to the weight matrix and bias vector). This model structure enables the differentiation of microbial taxa by employing a slab component to probabilistically highlight taxa with a potentially significant impact, while the spike component minimizes the influence of taxa with lesser or negligible impact on model performance (Fig. 1) [37–39].

In the second part of VBayesMM, the decoder segment generates conditional distributions for predicting metabolite abundances from the latent representation estimated during the encoding phase (Fig. 1). Like the MMvec approach, the metabolite abundances are modeled using a multinomial distribution. This architecture is specifically designed to systematically associate microbial taxa with metabolite abundances on a probabilistic basis, thereby enabling more accurate and interpretable predictions. Moreover, VBayesMM prioritizes a small subset of microbial species with the highest probability to be related to specific metabolite profiles (Fig. 1). These relationships are elucidated by the calculated co-occurrence probabilities. A more detailed description of the VBayesMM approach can be found in the Methods.

### 2.2 VBayesMM achieves accurate performance in large datasets

To evaluate the performance of VBayesMM, we applied it to four multiomics microbiome datasets derived from human and mouse samples (see Methods), which included thousands to tens of thousands of features such as microbiome species and metabolite abundances. We conducted a direct comparison using the default settings of the mmvec package in Python, and additionally employed the sparse partial least squares (sPLS) method from the mixOmics package in R, a tool designed for integrating multiple data types [18, 19]. The sPLS method was used with the default parameters of the *tune*.*spls* function from mixOmics package, allowing us to compare the performance of VBayesMM, MMvec, and sPLS across diverse datasets. The performance of each method was evaluated using the Symmetric Mean Absolute Percentage Error (SMAPE), a reliable metric for assessing their effectiveness across datasets of varying scales.

The SMAPE values computed across various datasets were used to assess the effectiveness of the VBayesMM method in comparison with MMvec and sPLS, as detailed in Figure 2 and Table S1. VBayesMM consistently achieved lower SMAPE values compared to MMvec and sPLS in all tested datasets. Specifically, in dataset A, VBayesMM had SMAPE values of 34.74% for the IHH case and 35.05% for air control, significantly lower than MMvec’s 47.59% and 48.17%, respectively. Both neural network-based methods performed better than the traditional sPLS approach, with SMAPE scores of 67.52% for the IHH case and 69.37% for air control. Supplementary Figure S1A supports these findings by illustrating the ELBO values for VBayesMM’s training and comparing the MAE values against MMvec during the validation phase. The graphical representations in Supplementary Figures S1Ac and S1Ad emphasize VBayesMM’s stability and lower MAE values *versus* the variable results of MMvec. This consistent performance underpins the potential for improved accuracy and reliability in multiomics data analysis. In dataset B, where the number of metabolite abundances was notably higher at 2,257 compared to 382 in dataset A, and the number of taxonomic units was substantially lower at 913 versus 4,690 in dataset A, the performance metrics indicated an increase in SMAPE values. Specifically, VBayesMM achieved a SMAPE of 55.07%, lower than MMvec’s score of 60.77% and sPLS’s score of 78.32%. This result suggests that while the effectiveness of VBayesMM diminishes with an increase in metabolite abundances relative to microbiome taxonomic units, it still outperforms the other methods, indicating a potential area for further optimization or adjustment in the algorithm’s application to similar scenarios. These comparisons highlight VBayesMM’s overall superior performance and stability in multi-omics data integration.

**Fig. 2:**
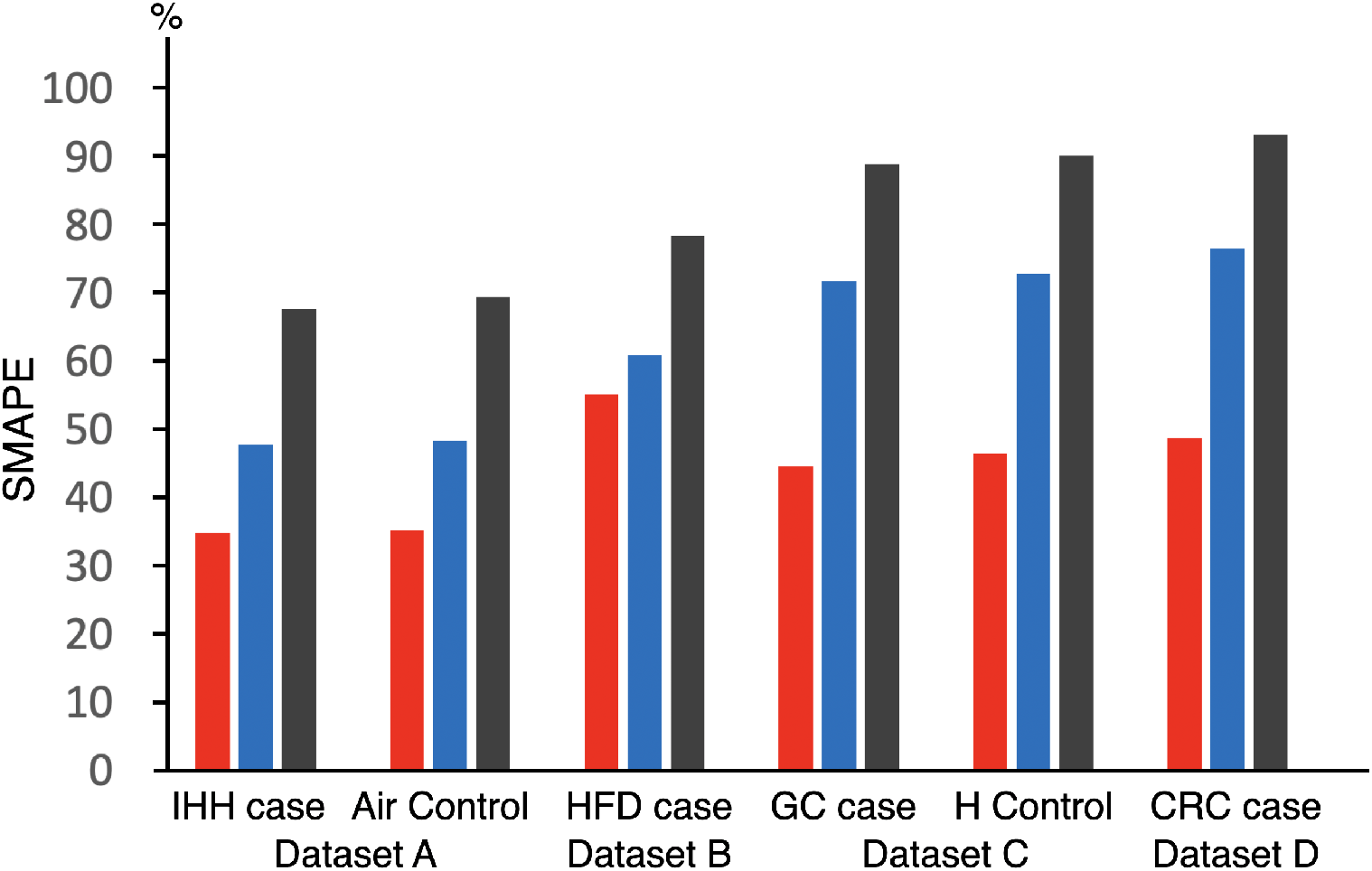
Symmetric Mean Absolute Percentage Error (SMAPE) values are computed for three approaches on real datasets. The VBayesMM approach is denoted by red color, mmvec by blue, and mixOmics by black. Dataset A includes cases of intermittent hypoxia and hypercapnia (IHH) and air control. Dataset B includes a high-fat diet (HFD) case. Dataset C comprises a gastric cancer (GC) case and a healthy (H) control. Dataset D contains a colorectal cancer (CRC) case. Note: All algorithms were run on a personal computer (Intel® Xeon® Gold 6230 Processor 2.10 GHz × 2, 40 cores, 2 threads per core) under Ubuntu 22.04.4 LTS.

To evaluate the efficacy of VBayesMM across different metagenomic sequencing methods, we used integrated shotgun metagenomic sequencing and mass spectrometry datasets. These datasets are known for their higher complexity and dimensionality compared to 16S datasets [53, 54]. Specifically, dataset C showcases an ultra-high dimensional context with 48,243 taxonomic units, far exceeding those in datasets A and B. However, the number of metabolite abundances (183) is smaller than in datasets A (382) and B (2,257). Figure 2 and Supplementary Table S1 show higher SMAPE values in dataset C compared to datasets A and B. Nonetheless, VBayesMM demonstrates a performance improvement in the GC case and health control, with SMAPE values of 44.42% and 46.31%, respectively. This represents a substantial enhancement of performance over the MMvec results, which achieved SMAPE values of 71.58% and 72.79%, respectively. Additionally, Supplementary Figures S1Cc and S1Cd illustrate VBayesMM’s superior performance in both GC case and health control, displaying consistently lower and more stable MAE values, in contrast to the fluctuation observed with the MMvec approach.

Despite dataset D featuring the largest number of taxonomic units among the evaluated datasets, totaling 57,702, it had a relatively small number of metabolite abundances at just 169. Nevertheless, the VBayesMM approach demonstrated improved performance for this dataset, achieving a SMAPE value of 48.64%, markedly better than the 76.45% for the MMvec. Supplementary Figure S1Db illustrates this improvement, showing that the MAE value for the VBayesMM was not only significantly lower but also more stable than the MAE value for the MMvec. Figure 2 shows that the neural network methods again performed better than the traditional method sPLS, particularly in the integrative analysis of shotgun metagenomics and multi-omics datasets. These results suggest that VBayesMM enhanced accuracy and stability in analyzing ultra-high dimensional datasets compared to traditional and alternative neural network approaches.

### 2.3 VBayesMM identifies a core set of features for microbial species

To evaluate the effectiveness of VBayesMM in identifying keystone microbial species that improve the accuracy of co-occurrence probability estimates between microbes and metabolites, we used the posterior distributions of the latent microbiome matrix **U** and microbial species selection (denoted as ***γ***). These distributions are shown in Figures 3a and 3c for dataset A, with Supplementary Figure S2a providing additional insights for dataset B.

**Fig. 3:**
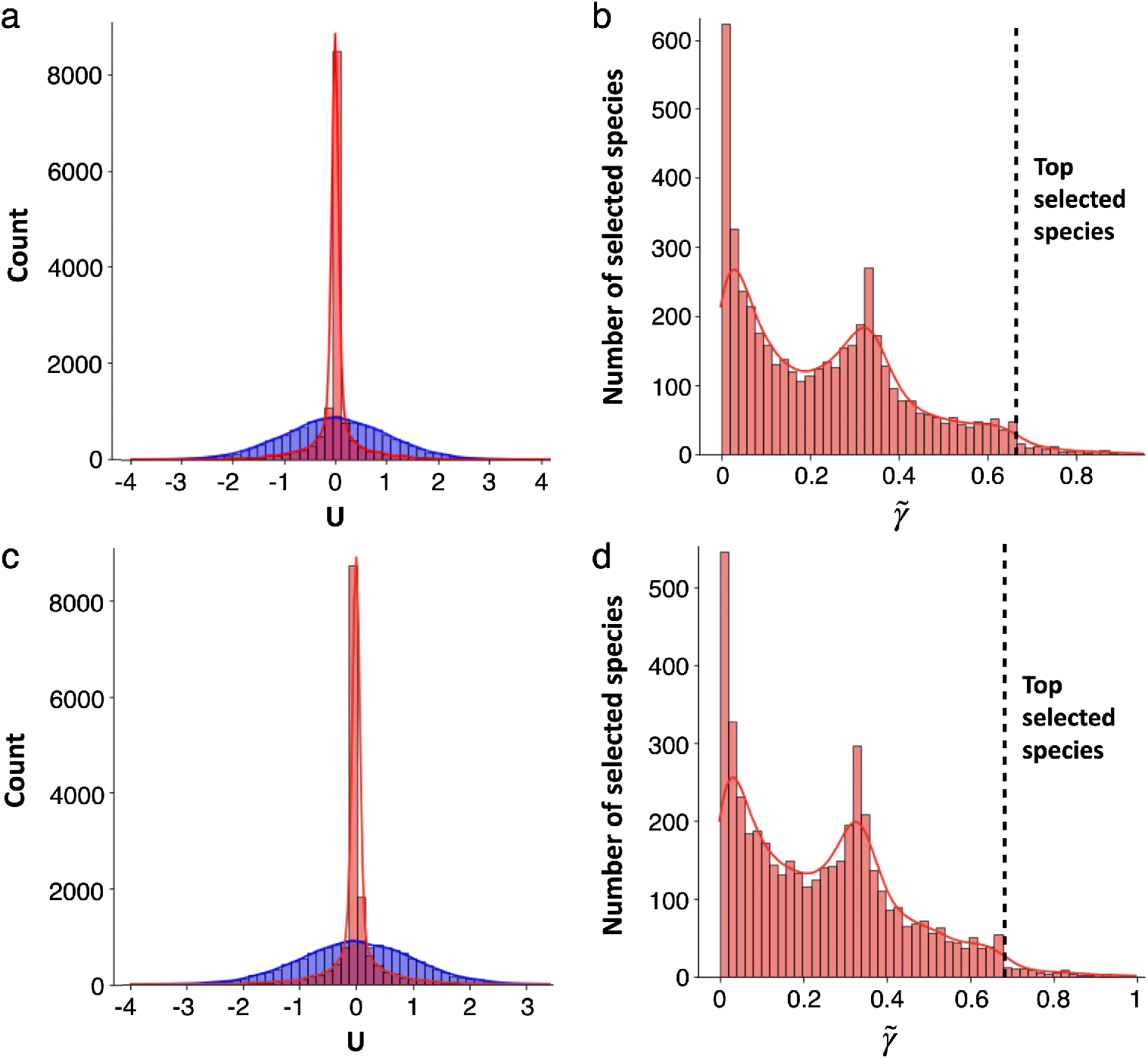
Histogram of the posterior probability distribution **U** and the average of 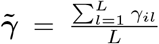 in dataset A. The VBayesMM and mmvec approaches are represented in red and blue respectively. The dashed lines are bound to select microbiome species. a and b. intermittent hypoxia and hypercapnia (IHH) cases group; c and d. controls group.

Specifically, VBayesMM is better at removing the non-informative microbial species because it takes advantage of the spike-and-slab distribution. Unlike the normal distribution used in MMvec, it is characterized by a sharp peak at origin, which leads to most of the variable weights to be set to zero. The spike component of the distribution aggressively enforces sparsity, which is highly beneficial in this case as it aligns with the baseline assumption that most microbial species will not significantly contribute to the model. Conversely, the slab component of the distribution, represented by the distribution’s tails, retains a subset of microbial species deemed potentially important, allowing their contributions to be accurately estimated without being affected by substantial shrinkage.

In contrast, the normal distribution used in the MMvec approach, with its broader spread, implies a more uniform assignment of variable weights, making non-zero weights more likely. This can lead to noise and potentially irrelevant microbial species being included in the model. This advantage of the spike-and-slab approach is demonstrated in comparisons that were done for these two strategies, where we have shown improvements in distinguishing influential from non-influential species, and, consequently, relevant co-occurrence probability estimates.

To illustrate how VBayesMM identifies a minimal core set of microbial species, we used the histograms showing the average posterior probability *γ*_*il*_ across the latent dimension *l*^*th*^ within dataset A, expressed as 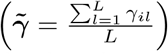, as shown in Figures 3b and 3d. Notably, the probability of 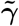 peaks sharply near 0, indicating that a substantial number of species have a very low probability of being influential, and therefore are not selected. This strict enforcement of sparsity by VBayesMM systematically excludes species with small likelihood contributions, simplifying the model and facilitating identification of species with more significant impacts.

Another secondary peak at 0.35 suggests a threshold for potentially influential species, with probability decreasing markedly after this point, indicating fewer species passing the threshold beyond this level. The selection threshold is indicated by a dotted line, and any species above it are considered highly influential. For example, we identified the top 50 microbial species with 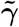 values over 0.7 for both IHH cases and control. In addition, Supplementary Figures S2a and b show the posterior distributions of **U** and the probability of 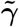 in dataset B. Here, we selected the top 50 microbial species with 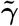 values exceeding 0.7 in the high-fat diet (HFD) samples. The selection is notably non-uniform, prioritizing species with higher 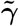 values. These figures illustrate the ways in which the proposed probability-based selection process can facilitate better understanding of how key microbial species contribute to the metabolite composition of the samples.

To confirm the biological significance of microbial species identified by VBayesMM in obstructive sleep apnea (OSA) study, we mapped the top 50 species associated with IHH cases and controls in dataset A onto the 16S phylogenetic tree, as shown in Figures 4a and 4b. The analysis highlighted species from the *Lachnospiraceae, Oscillospiraceae* and *Ruminococcaceae* families within the *Bacillota* phylum as particularly relevant to IHH case. Similar observations were reported in previous works [50]. Additionally, several studies on OSA have reported an increased prevalence of *Lachnospiraceae* and *Ruminococcaceae*, which are recognized for their fermentative abilities and associations with systemic inflammation, adipose tissue changes, and shifts in insulin sensitivity. These changes in gut microbiota may affect metabolic health through mechanisms such as disruption of the colonic epithelial barrier [55, 56].

**Fig. 4:**
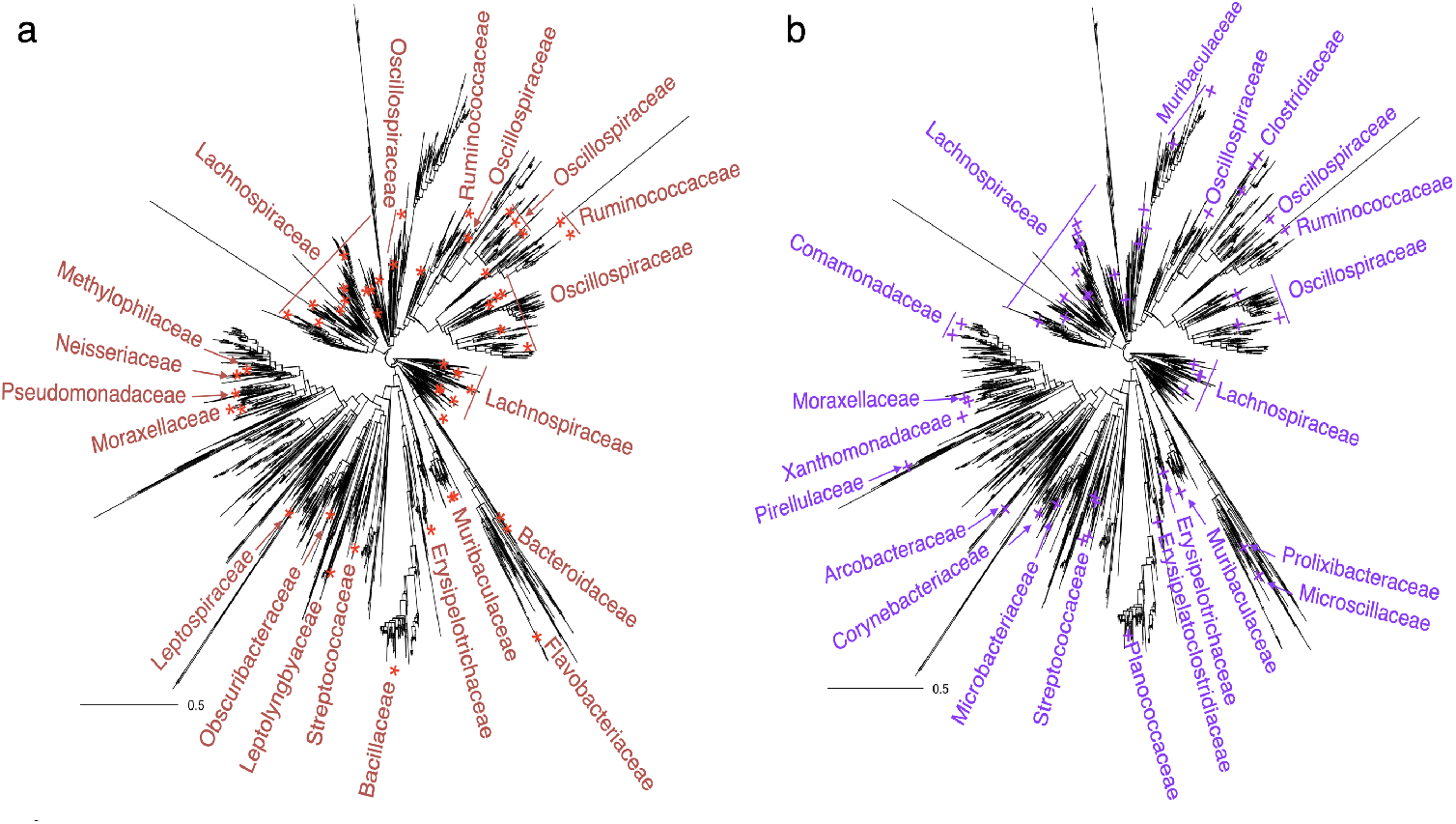
Microbial species selected using the VBayesMM approach and mapped on the phylogenetic tree based on 16S rRNA gene sequences for dataset A. a. intermittent hypoxia and hypercapnia (IHH) cases group in red color; b. controls group in purple color.

Additionally, families from the *Pseudomonadota* phylum, including *Methylophilaceae, Neisseriaceae, Pseudomonadaceae*, and *Moraxellaceae* are exclusively linked to IHH exposure (Figure 4a). Figure 4b illustrates that other families within the *Pseudomonadota* phylum, such as *Comamonadaceae* and *Xanthomonadaceae* families, are more abundant in the control group. These observations align with prior studies [57–60]. For example, members of *Neisseriaceae* family are known for their ability to reduce nitrate to nitrite in the oral cavity, which can be absorbed and converted into nitric oxide, a blood pressure regulator. Previous studies have shown significant enrichment of these bacteria in the hypertensive group [57, 58], suggesting a link to inflammation and certain cardiovascular conditions, and highlighting their potential impact on systemic health through their metabolic functions.

Previous research has consistently emphasized the substantial impact of the *Gammaproteobacteria* class, including families such as *Pseudomonadaceae* and *Moraxellaceae*, on obstructive sleep apnea [59, 60]. This bacterial class is closely associated with inflammatory processes within the human body, comprising multiple species directly involved in various pathological conditions. Notably, an increase in the *Gammaproteobacteria* class within the microbiome strongly correlates with elevated levels of pro-inflammatory cytokines, such as interleukin-6 [59]. This relationship highlights the critical role these bacteria play in exacerbating the inflammatory conditions observed in OSA, suggesting that their presence may significantly influence the severity and progression of the disease.

In the next application case, we applied the VBayesMM approach to determine the association of various microbial compositions with a high-fat diet (HFD) in dataset B. This analysis has identified top 50 critical microbial species displayed on the 16S phylogenetic tree in Supplementary Figure S3. This included keystone species from the *Lachnospiraceae, Oscillospiraceae* and *Ruminococcaceae* families within the *Bacillota* phylum, as well as *Muribaculaceae, Rikenellaceae*, and *Marinifilaceae* families from the *Bacteroidota* phylum. Recent studies indicate that high-fat diets induce a significant shift in gut microbiota, notably marked by an increased presence of *Lachnospiraceae*. This shift underscores their pivotal role in energy metabolism, which may be driving the increased adiposity and obesity observed in experimental models, while also emphasizing their substantial impact on metabolic health [61–63]. The pronounced abundance of *Lachnospiraceae* in response to high-fat dietary patterns strongly correlates with greater risks of metabolic disorders and diabetes, suggesting that these bacteria may play a crucial role in exacerbating the metabolic patterns associated with obesity [61]. Emerging evidence highlights the substantial role of dietary fats in modulating gut microbiota including its downstream effects on overall metabolic health and disease progression.

### 2.4 VBayesMM prioritizes the co-occurrence probabilities between the most informative microbial species and metabolite abundances

To evaluate the probabilistic relationships between keystone microbial species and metabolite abundances in IHH case and control groups within dataset A, we analyzed the co-occurrence probabilities between microbes and metabolites using heatmaps (Figures 5a and 5b). These heatmaps illustrate the strength of probabilistic associations between various microbial species and specific metabolites. For example, Figure 5a highlights that microbial families such as *Oscillospiraceae, Muribaculaceae*, and *Methylophilaceae* are prevalent in environments with high concentrations of cholic acid, indicating robust cooccurrence probabilities under IHH exposure. Similarly, a prominent presence of *Lachnospiraceae* in environments rich in chenodeoxycholic acid correlates with their high co-occurrence probabilities. These findings are consistent with previous research, and also expand our understanding of microbial interactions with metabolites in OSA scenarios [50, 64–66].

**Fig. 5:**
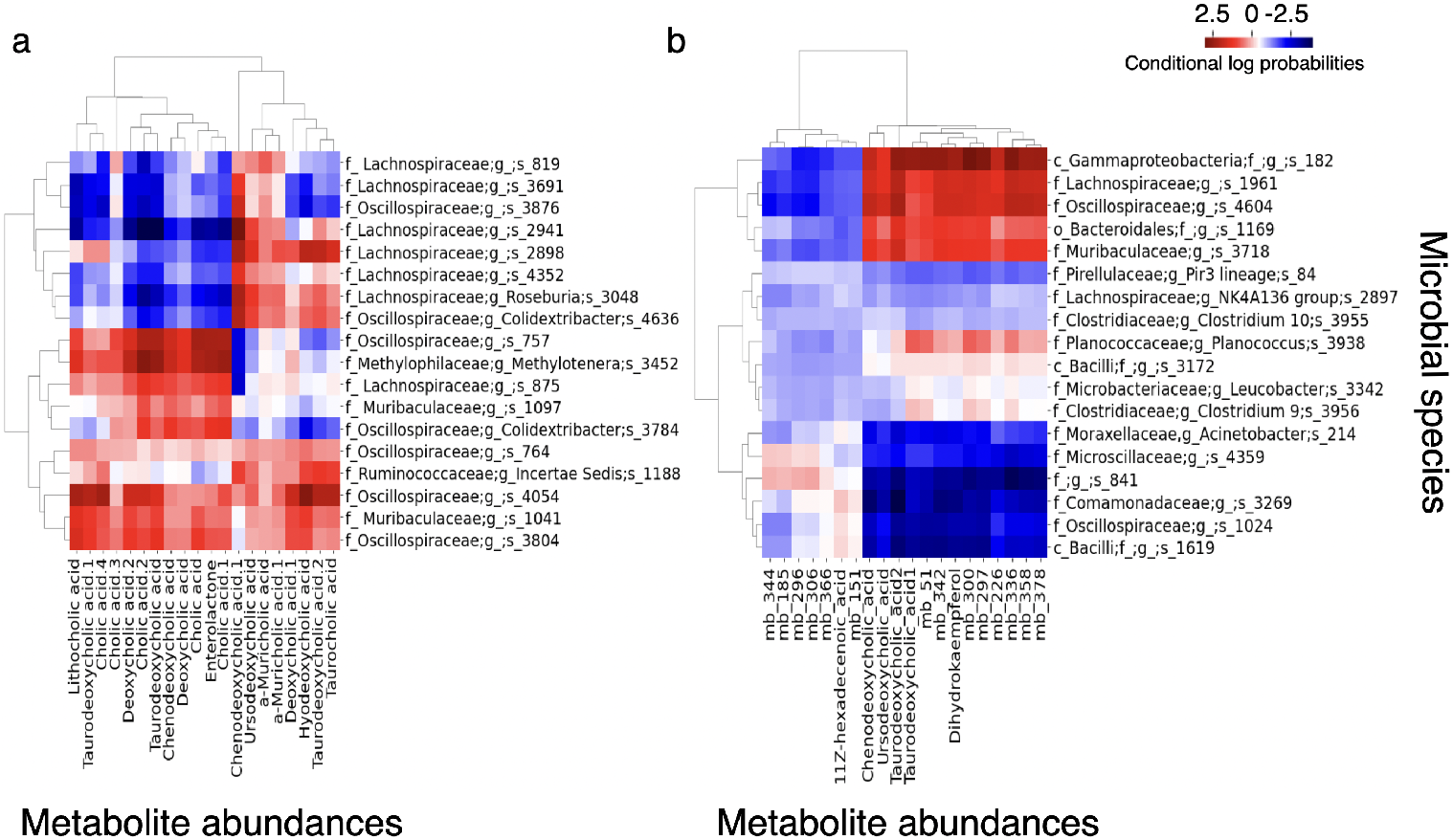
Heat map of the estimated conditional log probabilities of VBayesMM for the selected microbial species and metabolite abundances in dataset A. Individual metabolites and microbiomes were hierarchically clustered (Ward’s method) using Euclidean distance. a. intermittent hypoxia and hypercapnia (IHH) cases group; b. controls group. Note: f denotes family; g denotes genus; s denotes species; mb denotes metabolite.

In particular, interaction between the *Lachnospiraceae* family and chenodeoxycholic acid appears to be a key factor in the metabolic implications in OSA. The *Lachnospiraceae*, known for their role in converting primary to secondary bile acids, may experience functional changes due to OSA-related disruptions in gut barrier function. Such disruptions could contribute to systemic inflammation and affect the microbial balance in the gut, potentially impacting bile acid metabolism [64, 65]. Chenodeoxycholic acid, a primary bile acid, plays roles not only in bile production but also as a signaling molecule [65]. It is crucial in pathways linked to inflammation and oxygen sensing, which are highly relevant to the pathophysiology of OSA. The fluctuating oxygen levels characteristic of OSA may trigger metabolic disturbances with chenodeoxycholic acid modulating the body’s response to intermittent hypoxia. These interactions underscore the importance of understanding biochemical dynamics for better management of OSA and its metabolic implications.

Further analysis using the VBayesMM approach on dataset B (Supplementary Figure S4) focused on selected probabilistic relationships between microbes and metabolites in a high-fat diet study. This analysis revealed an association between the *Lachnospiraceae* family and hyodeoxycholic acid (HDCA) [67, 68]. As was noted earlier, the *Lachnospiraceae* family plays an important part in bile acid metabolism, by converting primary bile acids into secondary forms. HDCA is one such secondary bile acid and is synthesized from chenodeoxycholic acid via bacterial enzymatic activity within the gut. Under a high-fat diet, the abundance of the *Lachnospiraceae* family increases – a shift that closely correlates with variations in HDCA levels [68]. This relationship emphasizes the transformative role of *Lachnospiraceae* in modulating bile acid profiles, which is pivotal to understanding dietary impacts on metabolic health. These findings offer insights into how dietary fats influence microbial dynamics and their subsequent effects on host metabolism.

## 3 Discussion

Accurate identification of core taxonomic units (or microbial species) within the microbiome is essential for enhancing our understanding of how it relates to other omics data, most importantly from the perspective of microbe–metabolite interactions. The approach proposed in this paper aims to improve quality of such inferences and open up new research opportunities in exploring the role microbiome plays across diverse fields such as human disease, precision medicine, and environmental studies.

The VBayesMM method represents a significant advancement in microbiome multiomics analysis, addressing the challenges posed by high dimensionality of integrated datasets, which often led to scalability issues, reduced performance, and interpretation challenges when analyzed with previous methods. VBayesMM employs a spike-and-slab prior regularization on the weight matrix and bias vector of the encoder neural network, allowing for the probabilistic identification of a limited number of microbial species (or taxonomic units) statistically found to improve model performance. At the same time, variational inference strategy is used to address the computational bottlenecks associated with traditional Bayesian sampling algorithms to ensure both timely execution and improved predictive accuracy. The scalability of VBayesMM is demonstrated by successfully applying it to the extensive microbiome datasets derived from shotgun metagenomic sequencing [51, 52].

Recent studies support the use of neural networks for classifying microbial features and predicting host phenotypes solely from shotgun metagenomics datasets [69, 70]. VBayesMM facilitates the discovery of probabilistic relationships between metabolites and the microbiome’s community structure and function, offering valuable insights for understanding their impact on human health and disease. Additionally, VBayesMM incorporates a Bayesian neural network that quantifies uncertainties, enabling detailed probabilistic interpretations of predicted outcomes. This capability allows for the estimation of variational posterior probabilities across microbes and metabolites. The interpretability and scalability of VBayesMM facilitate its application across diverse datasets, make it a valuable tool for integrating various microbiome datasets and supporting multiomic analysis in complex biological systems.

While we proposed several solutions to address critical challenges in microbiome multiomics analysis, VBayesMM is not without limitations. The model is primarily designed to optimize the contributions of microbiome data, which is optimal for its primary analytical objectives. However, its stability and performance can be compromised when the number of metabolite abundances significantly exceeds the number of microbial species, such as in the dataset from the high-fat diet study (dataset B). This discrepancy can affect the stability and accuracy of the VBayesMM. Future developments can explore further improvements necessary to tackle the dimensionality challenges presented by mass spectrometry metabolomics datasets. Additionally, the integration of more comprehensive omics analyses is become increasingly relevant, for example as those combining metatranscriptomics and metabolomics or metaproteomics and metabolomics [71–73]. Such integrative approaches are crucial for uncovering the complex interactions between microbial communities and the human host. To address these emerging needs, further enhancements to VBayesMM should focus on improving data integration across various omics technologies to deliver even more accurate identification of associations between microbes and specific nutrients or medications. These developments are expected to enhance the analytical capabilities of VBayesMM and expand its utility in diverse biological datasets, potentially enriching our understanding of microbiome-related health dynamics.

## 4 Methods

### 4.1 Data acquisition

We used microbiome-metabolite data from previously published datasets composed of both mouse and human samples. Dataset A is from an experiment that studied the obstructive sleep apnea (OSA) in mice, including 92 samples for intermittent hypoxia and hypercapnia (IHH) and 90 samples for control conditions, along with 4,690 taxonomic units and 382 metabolite abundances [50]. The metabolomics detection and annotations were performed using the Global Natural Product Social Molecular Networking platform (GNPS) [74]. This analysis identified a number microbe-dependent metabolites such as bile acids (chenodeoxycholic acid, cholic acid), phytoestrogens (enterodiol, enterolactone), and fatty acids (elaidic acid, phytomonic acid) that had different levels in between the IHH and control groups [50]. Dataset B, from a study on effects of high-fat diet (HFD) in a murine model, has 438 samples, 913 taxonomic units, and 2,257 metabolite abundances [16]. The GNPS facilitated the determination of metabolites such as secondary bile acids, primary bile acids, soyasaponins, and peptides, which are microbially produced. Datasets A and B were analyzed using 16S rRNA gene sequencing-based microbiome and liquid chromatography-tandem mass spectrometry (LC-MS/MS)-based metabolome datasets.

Dataset C was generated from profiling the fecal microbiome and metabolites in gastric cancer (GC) patients for 42 samples of gastrectomy cases and 54 samples of health controls, which include 48,243 taxonomic units and 183 metabolite abundances [52, 75]. Annotation of these metabolites was summarized into Kyoto Encyclopedia of Genes and Genomes (KEGG) [76]. This annotation process helped in the identification of key microbe-dependent metabolites such as glycocholate, taurine, and cholate, which were found to be significantly enriched in the gastrectomy group compared to controls [52]. Dataset D focuses on the fecal microbiome and metabolites in colorectal cancer (CRC) patients from stage 0 to stage 4, with 150 samples featuring 57,702 taxonomic units and 169 metabolite abundances [51, 75]. The KEGG annotation identified bacterial metabolites such as bile acids and short-chain fatty acids, which have established associations with the CRC stages. Datasets C and D have whole-genome shotgun sequencing (WGS) microbiome profiling and capillary electrophoresis time-of-flight mass spectrometry (CE-TOFMS) for metabolomics.

### 4.2 The architecture of the MMvec neural network

First, we provide a concise overview of the microbe–metabolite vectors (MMvec) approach, which estimates co-occurrence probabilities to elucidate the relationships between microbiomes and metabolites [16]. This method enables the determination of metabolite abundance data via conditional distributions based on microbiome abundances, utilizing the multinomial distribution. The fundamental concepts of the MMvec approach are as follows: Given the paired microbiome-metabolome dataset **D** = {**X, Y**}, which includes K samples, N taxonomic units (or microbial species), and M metabolite abundances, we denote the observations for the *i*^*th*^ taxonomic unit and the *j*^*th*^ metabolite in the *k*^*th*^ sample as *X*_*ki*_ and *Y*_*kj*_ respectively, where *k* ∈ {1, …, *K*}, *i* ∈ {1, …, *N*}, and *j* ∈ {1, …, *M*}. The MMvec approach maps taxonomic units and metabolite abundances into a latent low-dimensional space of dimension L. Here, each *i*^*th*^ taxonomic unit is represented by the vector **u**_*i*_ ∈ ℝ^*L*^, and each *j*^*th*^ metabolite abundance by the vector **v**_*j*_ ∈ ℝ^*L*^. These latent vectors are drawn from a prior normal distribution with a mean of 0 and a diagonal covariance matrix I, where the variances for the taxonomic units and metabolites are *σ*_*i*_ and *σ*_*j*_ respectively:

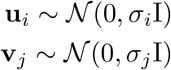

For a given microbial sample **X**_*k*_ = (*X*_*k*1_, …, *X*_*kN*_), each *i*^*th*^ taxonomic unit (or microbial species) is sampled from a categorical distribution, denoted as *i* ∼ Categorical(**X**_*k*_). Given sample *k* and the total metabolite abundances across sample *n*, the MMvec models the metabolite abundances **Y**_*k*_ = (*Y*_*k*1_, …, *Y*_*kM*_) as being drawn from a multinomial distribution as follows:

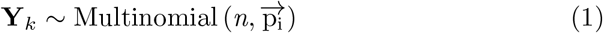

where 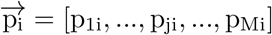 is the conditional probability distribution for observing *j*^*th*^ metabolite abundance given that *i*^*th*^ taxonomic unit is observed. This is defined as follows:

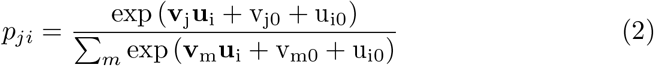

where *v*_*j*0_ and *u*_*i*0_ are biases used to accurately approximate the conditional distribution of microbiome-metabolite relationships. To refine the estimates, maximum a posteriori (MAP) estimation is used to optimize the objective function with respect to the matrices **U** and **V** [16]. This optimization is done using the ADAM algorithm [77]:

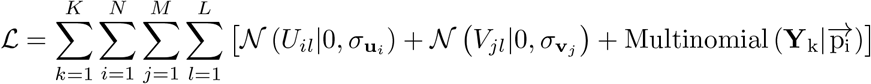

where **U** = [0, **u**_0_, **u**_1_, …, **u**_*N*_] ∈ ℝ^*N ×L*^ denotes the embedding for N taxonomic units (or microbial species) and **V** = [**v**_0_, 0, **v**_1_, …, **v**_*M*_] ∈ ℝ^*M −*1*×L*^ denotes the embedding for M metabolite abundances.

### 4.3 Variational Bayesian microbiome multiomics (VBayesMM) approach

In the MMvec approach, all microbial species are initially assumed to equally contribute to the prediction of the complete metabolite abundances profile. However, this assumption is often not appropriate in microbiome-metabolite research [78, 79], as a large number of microbial species — potentially amounting to tens of thousands of taxonomic units — may have negligible relevance. In reality, only a limited number of microbial species are involved in relationships with metabolites, and these relationships impact the estimations of the co-occurrence probabilities between microbes and metabolites. To address this issue, we propose incorporating a spike-and-slab prior, a method from Bayesian analysis [37–39, 80], into the latent microbiome matrix **U** of Bayesian neural network. This approach aims to prioritize the core set of microbial taxonomic units within the MMvec neural network. Consequently, the prior distribution of *i*^*th*^ taxonomic unit **u**_*i*_ can be defined as follows [37–39]:

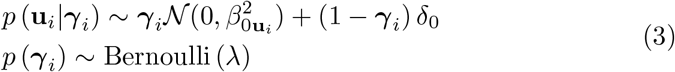

where ***γ***_*i*_ ∈ ℝ^*L*^ is a binary vector, such as ***γ***_*i*_ = 1 indicates that the *i*^*th*^ taxonomic unit is important for estimating the conditional distribution of *j*^*th*^ metabolite abundance in equation 2 and ***γ***_*i*_ = 0 denotes that the *i*^*th*^ taxonomic unit is unimportant for learning the co-occurrence probabilities. *δ*_0_ denotes the Dirac spike concentrated at zero and *λ* is the hyperparameter of prior Bernoulli distribution. The prior distribution of *j*^*th*^ metabolite abundance **v**_*j*_ follows normal distribution 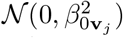.

We propose the Variational Bayesian (VB) method [37–39, 81] to perform efficient approximate posterior inference, which is necessary for learning the co-occurrence probabilities between the microbiome and metabolome. Utilizing the observed omics datasets, denoted as **D**, our framework uses several key variables: **Ξ** which includes the embedding matrix for microbial species **U**, a binary matrix for taxonomic unit selection ***γ***, and the embedding matrix for metabolite abundances **V**. To manage the computational complexity and intractability of the true posterior distribution *p*(**Ξ** |**D**), we introduce a variational distribution *q*(**Ξ** | Θ), characterized by the hyperparameter Θ. This approach allows for a tractable and effective approximation of the posterior distribution. In VBayesMM, these variational distributions are defined as follows:

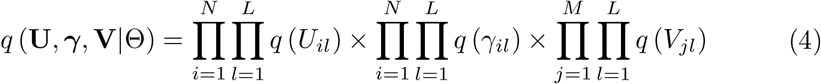

where

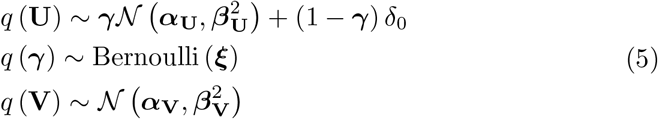

In equation 5, the variational parameter Θ comprises {***α***_**U**_, ***β***_**U**_, ***ξ, α***_**V**_, ***β***_**V**_} The distributions of the exponential families were selected for the variational distributions to guarantee a feasible computation of the expectations. The log marginal probability log (p (**D**)), which is known as the evidence of **D**, is defined as follows:

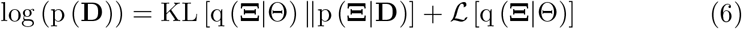

where the first term of equation 6 is the KL divergence to measure the similarity between *q* (**Ξ** |Θ) and *p* (**Ξ** |**D**). Since KL [q (**Ξ** |Θ) ∥ p (**Ξ** |**D**)] ≥0, the variational Bayesian approach optimizes the second term of equation 6, which is called Evidence Lower Bound (ELBO), defined as follows:

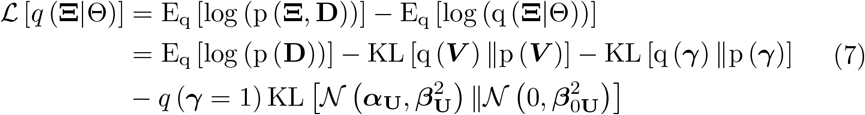

In equation 7, the variational parameters for the normal variables **U** and **V** are reparameterized using the expression ***α*** + ***βϵ***, where ***ϵ*** follows a standard normal distribution, 𝒩 (0, 1) [81]. Since the variable ***γ***, which represents microbial selection, is discrete, the continuous reparameterization trick is not applicable. To address this, we adopt the Gumbel-softmax approximation [37–39], which facilitates the reparameterization of categorical variables as follows:

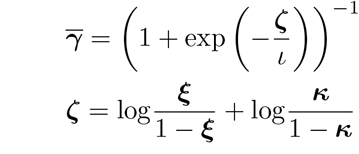

where ***κ*** ∼ *U* (0, 1) and *?*, which represents the temperature parameter, is set to a minimum of 0.5 to ensure numerical stability [37–39]. With all variational distributions *q*(**Ξ**|Θ) reparameterized into differentiable forms, we utilize the stochastic gradient approach [77] to optimize the ELBO ℒ [*q* (**Ξ**|Θ)]. The mathematical details of the gradient of ELBO ∇_(Θ)_ ℒ [*q* (**Ξ** |Θ)] with respect to variational parameters are provided in the Supplementary Material. The complete procedure of VBayesMM is defined as follows:

#### Algorithm 1

Variational Bayesian microbiome multiomics - VBayesMM

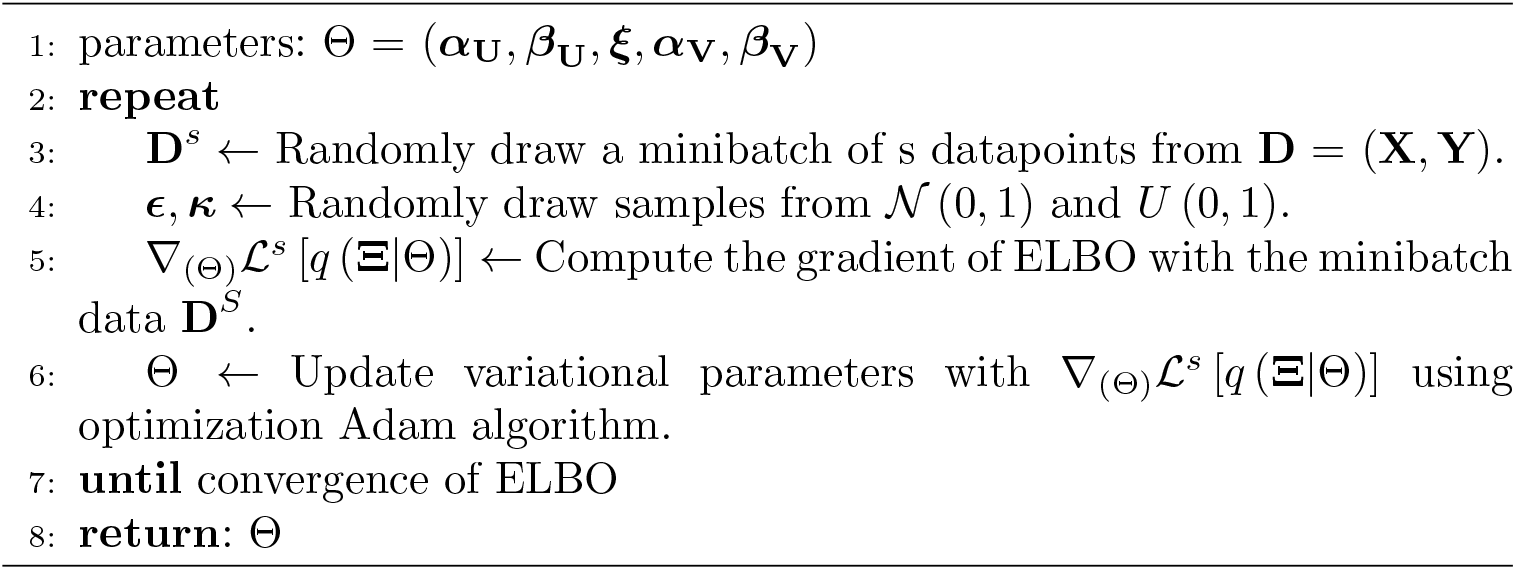

### 4.4 Criteria to evaluate the performance of the approaches

In both the MMvec and VBayesMM approaches, the data is actually counts of metabolite abundances, so the predictive performance on test samples can be determined using mean absolute error (MAE). The MAE is calculated as follows [16]:

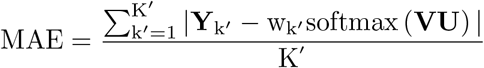

where *w*_*k*^*′*^_ denotes the total metabolite abundances in test sample *k*^*′*^ ∈ {1, …, *K*^*′*^}. For each microbial species, data is resampled from a categorical distribution based on the microbial composition of the testing sample **X**_*k*_*′*. To validate performance, 20% of the total samples across all four datasets were randomly set aside as test samples, while the remaining 80% were used for training.

Then, in order to assess the performance of these approaches across datasets, which often exhibit significant variations in the scale of observations, we use the symmetric mean absolute percentage error (SMAPE) as a robust measure of accuracy [82]. This metric provides a reliable comparison regardless of the dataset size or scale and is computed as follows:

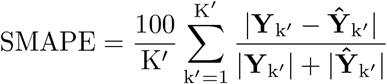

where Ŷ_*k*^*′*^_ is the predicted value and 0% ⩽ SMAPE ⩽ 100%.

### 4.5 Open-source software

The VBayesMM package has been implemented using Tensorflow, the same framework used to develop MMvec, with an additional version available in PyTorch. This dual-framework compatibility allows users to select the most suitable version for their own system. The software accepts microbiome and metabolite count data in CSV, TSV, or BIOM formats. The primary outputs of both mmvec and VBayesMM include the microbiome matrix **U**, the metabolite matrix **V**, and a matrix of log conditional probabilities for the relationships between microbial species and metabolites. A unique feature of VBayesMM is the inclusion of a selection probabilities matrix for microbial taxonomic units, which is particularly valuable when dealing with large datasets. This matrix enables users to identify and focus on the most important taxonomic units that show significant relationships with metabolites, facilitating the construction of informative heatmaps. Additionally, the software provides the ELBO and MAE outputs to evaluate model performance.

Users can set the number of gradient descent iterations and batch sizes, which traditionally impact computational times. Our computations were conducted on an Intel® Xeon® Gold 6230 Processor, featuring 2.10 GHz × 2, with 40 cores and 2 threads per core, running Ubuntu 22.04.4 LTS. In scenarios involving dataset A, the runtime for VBayesMM, utilizing all 40 cores (equivalent to 80 logical processors), is typically around 1.5 hour to achieve model convergence. Dataset B, which contains 2,257 metabolite abundances, reached a convergence after running for about 8.5 hours. For datasets with a large number of microbial taxonomic units, such as dataset C, which includes 48,243 microbial taxa and 183 metabolite abundances, convergence was reached within approximately 2 days. Similarly, dataset D, which contains 57,702 microbial taxa and 169 metabolite abundances, requires about 5 days to converge. These timelines reflect the processing capacity and efficiency of our computational setup, allowing timely analysis despite the scale of the data. The details of computational time are provided in Supplementary Table S2. The software is available at https://github.com/tungtokyo1108/VBayesMM/.

In all our experiments, we set the number of latent dimensions to 3, the prior distributions of mean and standard deviation for the weight matrix and bias vector of microbial species **U** and metabolite abundances **V** that follow the standard normal distributions, 𝒩 (0, 1). For the Adam algorithm, the learning rate is 0.1, the exponential decay rate for the 1st-moment estimates is 0.8, and the exponential decay rate for the 2nd-moment estimates is 0.9 [16]. To address the selection of microbial species, we set the value of the temperature parameter *ι* to 0.5, the initial value of the hyperparameter *ξ* of Bernoulli prior distribution to *U* (0,1) and the value of the hyp erparameter *λ* to log 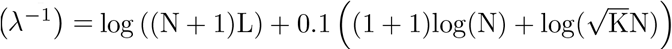 [37].

## Supplementary information

## Acknowledgments

This work was partly supported by JSPS KAKENHI Grant Number JP20H03240, JSPS KAKENHI Grant Number JP24K15175 and JST CREST Grant Number JPMJCR2231, Japan.

## Declarations

### Ethics approval and consent to participate

Not applicable

### Consent for publication

Not applicable.

