## Supplementary for "VBayesMM: Variational Bayesian neural network to prioritize important relationships of high-dimensional microbiome multiomics data"

### 1 Supplementary Figures

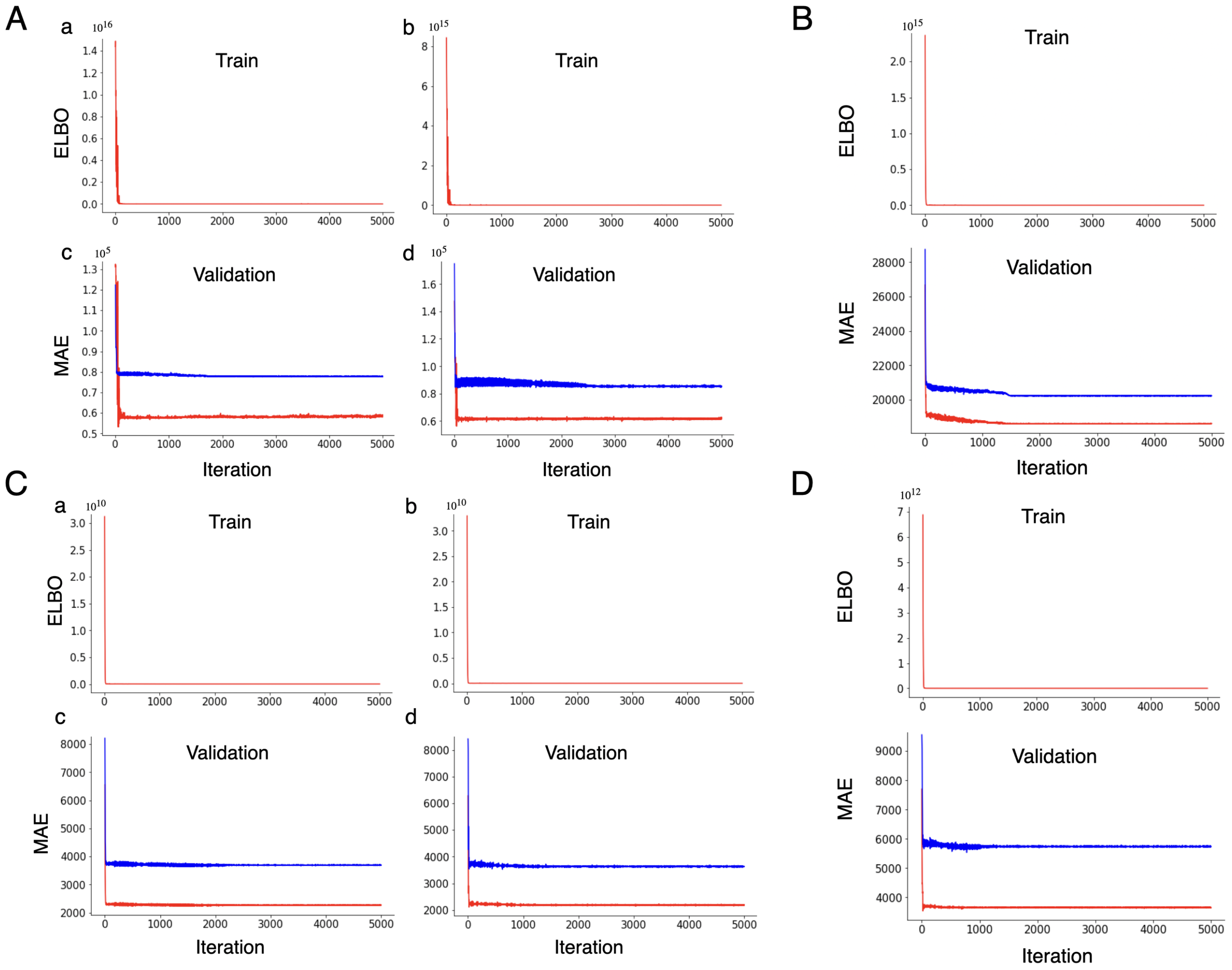


**Supplementary Figure S1**: The Evidence Lower Bound (ELBO) and Mean Absolute Error (MAE) values for the VBayesMM and mmvec approaches, represented in red and blue respectively. Panel A (Dataset A): Displays ELBO values for the intermittent hypoxia and hypercapnia (IHH) case and air control in parts (a) and (b), and MAE values in parts (c) and (d); Panel B (Dataset B): Shows the ELBO value in part (a) and the MAE value in part (b); Panel C (Dataset C): Features ELBO values for the gastric cancer (GC) case and healthy control in parts (a) and (b), with corresponding MAE values in parts (c) and (d); Panel D (Dataset D): Contains the ELBO value in part (a) and the MAE value in part (b).

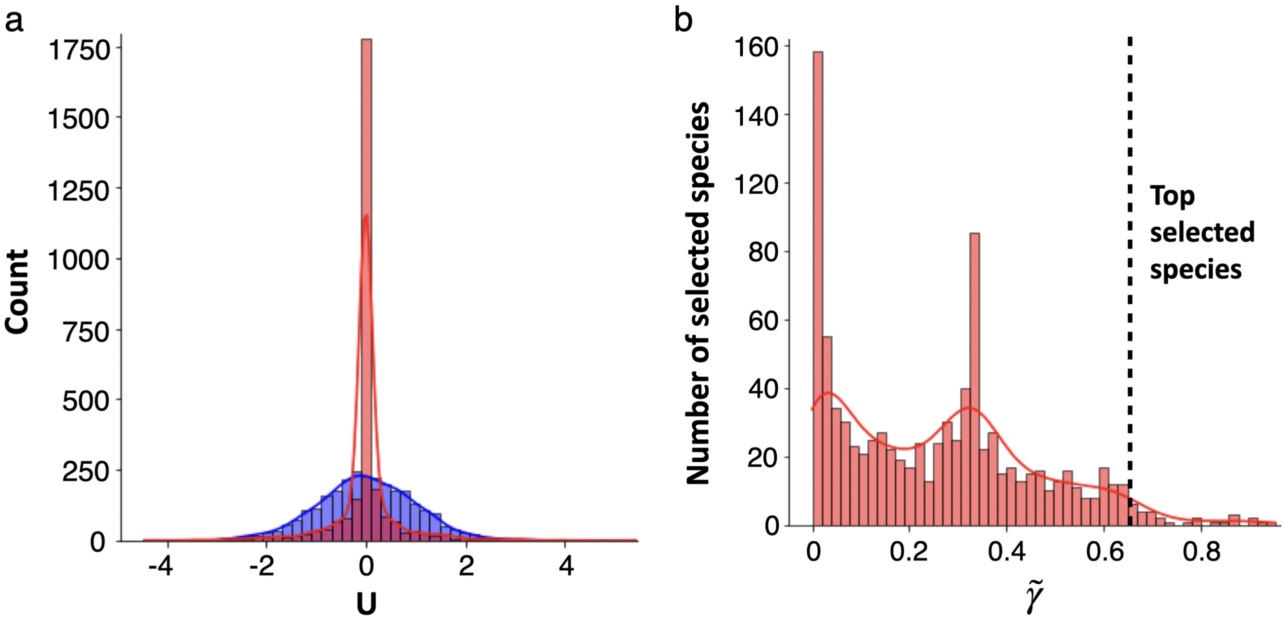


**Supplementary Figure S2**: Histogram of the posterior probability distribution $\mathbf{U}$ and the average of $\tilde{\gamma} = \frac{\sum_{l=1}^{L} \gamma_{il}}{L}$ in dataset B. The VBayesMM and mmvec approaches are represented in red and blue respectively. The dashed lines are bound to select microbiome species.

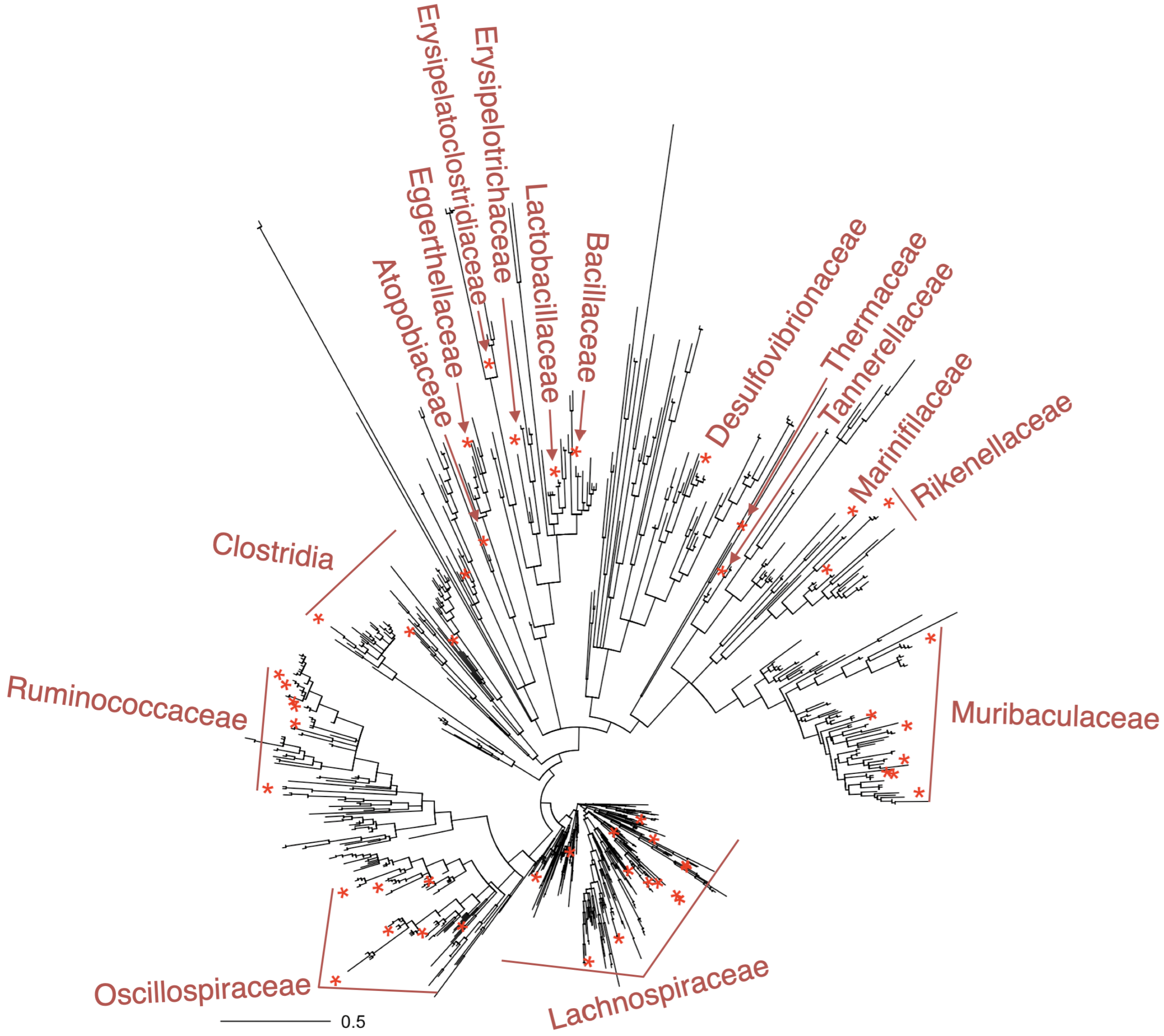


**Supplementary Figure S3**: Microbial species selected using the VBayesMM approach and mapped on the phylogenetic tree based on 16S rRNA gene sequences for dataset B.

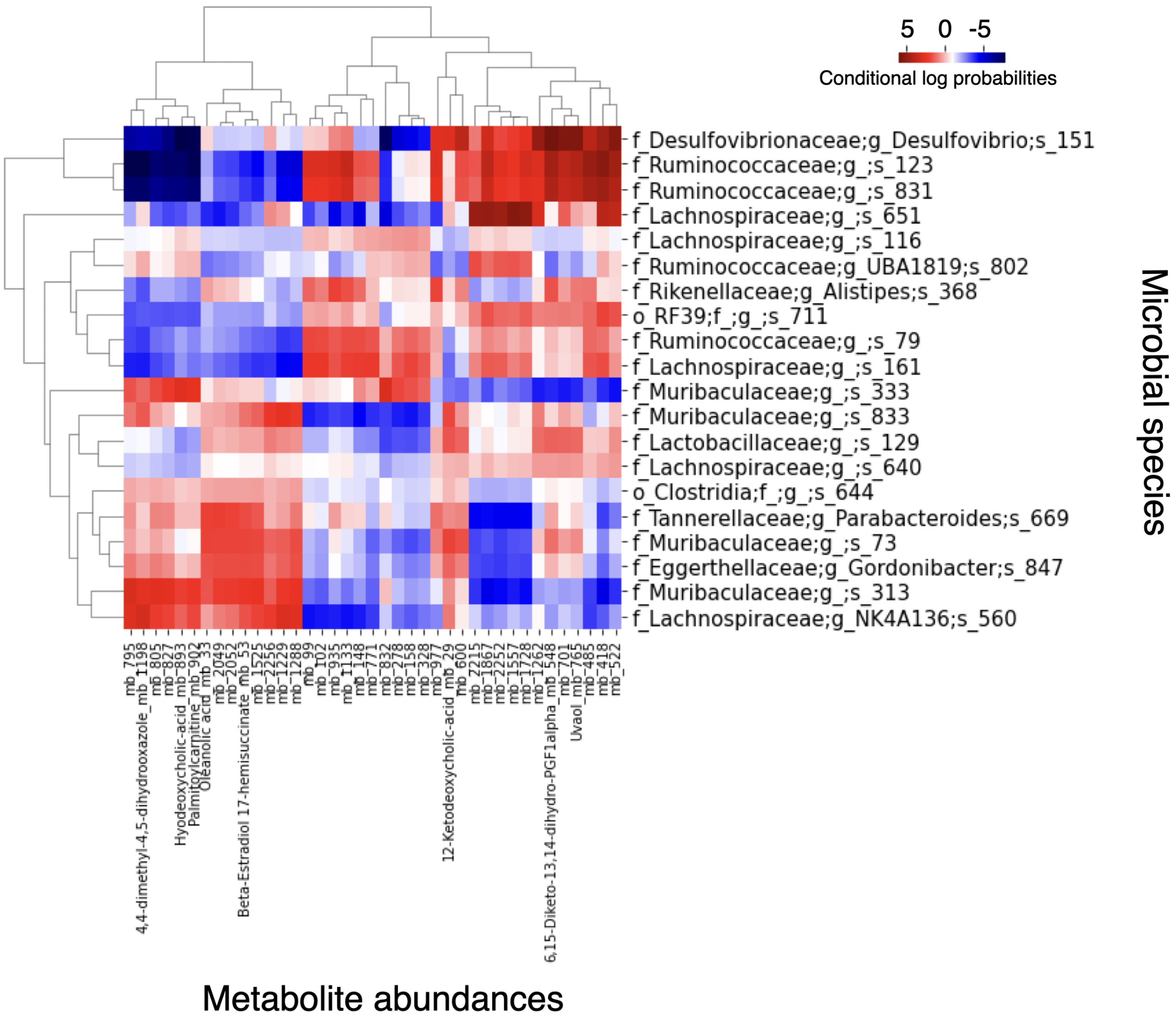


**Supplementary Figure S4**: Heat map of the estimated conditional log probabilities of VBayesMM for the selected microbial species and metabolite abundances in dataset B. Individual metabolites and microbiomes were hierarchically clustered (Ward's method) using Euclidean distance. Note: f denotes family; g denotes genus; s denotes species; mb denotes metabolite.

#### 2 Supplementary Tables

**Supplementary Table S1**: The SMAPE values of the three approaches on the real data sets.

| Dataset | Case–control | VBayesMM | MMvec | mixOmics |
|---|---|---|---|---|
| A | IHH case | 34.73 % | 47.59 % | 67.52 % |
| | Control | 35.06 % | 48.17 % | 69.37 % |
| B | HFD case | 55.07 % | 60.77 % | 78.32 % |
| C | GC case | 44.42 % | 71.58 % | 88.75 % |
| | Control | 46.31 % | 72.79 % | 90.03 % |
| D | CRC case | 48.64 % | 76.45 % | 93.12 % |

Note: SMAPE: symmetric mean absolute percentage error. IHH: intermittent hypoxia and hypercapnia. HFD: high-fat die. GC: gastric cancer. CRC: colorectal cancer. All algorithms were run on a personal computer (Intel® Xeon® Gold 6230 Processor 2.10 GHz $\times$ 2, 40 cores, 2 threads per core) under Ubuntu 22.04.4 LTS.

**Supplementary Table S2**: Running time of the three approaches on the real data sets.

| Dataset | Case–control | VBayesMM (Python) | MMvec (Python) | mixOmics (R) |
|---|---|---|---|---|
| A | IHH case | 1.52 h | 1.48 h | 2.63 h |
| | Control | 1.50 h | 1.45 h | 2.58 h |
| B | HFD case | 8.54 h | 8.28 h | 10.16 h |
| C | GC case | 48.66 h | 48.25 h | 51.19 h |
| | Control | 48.82 h | 48.43 h | 51.51 h |
| D | CRC case | 121.62 h | 120.97 h | 125.18 h |

Note: The VBayesMM and MMvec packages are implemented in parallel using Python. The mixOmics package is implemented in parallel using R.

### 3 Supplementary Methods

#### 3.1 Variational inference for Variational Bayesian microbiome multiomics (VBayesMM) approach

We expand specifically the Evidence Lower Bound (ELBO) equation as follows:

$$
\begin{aligned}
&\mathcal{L}\left[q\left(\boldsymbol{\Xi}|\Theta\right)\right] = \mathrm{E_q}\left[\log\left(\mathrm{p}\left(\boldsymbol{\Xi},\mathbf{D}\right)\right)\right] - \mathrm{E_q}\left[\log\left(\mathrm{q}\left(\boldsymbol{\Xi}|\Theta\right)\right)\right] \\
&= \mathrm{E_q}\left[\log\left(\mathrm{p}\left(\mathbf{D}\right)\right)\right] - \mathrm{KL}\left[\mathrm{q}\left(\boldsymbol{V}\right)\|\mathrm{p}\left(\boldsymbol{V}\right)\right] - \mathrm{KL}\left[\mathrm{q}\left(\boldsymbol{\gamma}\right)\|\mathrm{p}\left(\boldsymbol{\gamma}\right)\right] - \mathrm{q}\left(\boldsymbol{\gamma}=1\right)\mathrm{KL}\left[\mathcal{N}\left(\boldsymbol{\alpha}_{\mathbf{U}},\boldsymbol{\beta}^2_{\mathbf{U}}\right)\|\mathcal{N}\left(0,\boldsymbol{\beta}^2_{0\mathbf{U}}\right)\right] \\
&= \mathrm{E_q}\left[\log\left(\mathrm{p}\left(\mathbf{D}\right)\right)\right] - 0.5\sum_{\mathrm{j=1}}^{\mathrm{M}}\sum_{\mathrm{l=1}}^{\mathrm{L}}\left[\left(1+\log\left(\beta^2_{\mathrm{V_{jl}}}\right)-\alpha^2_{\mathrm{V_{jl}}}-\beta^2_{\mathrm{V_{jl}}}\right)\right] - 0.5\sum_{\mathrm{j=1}}^{\mathrm{M}}\left[\left(1+\log\left(\beta^2_{\mathrm{V_{j0}}}\right)-\alpha^2_{\mathrm{V_{j0}}}-\beta^2_{\mathrm{V_{j0}}}\right)\right] \\
&-\sum_{i=1}^{N}\sum_{l=1}^{L}\left[\left(\frac{1}{1+e^{-\xi_{U_{il}}}}\right)\times\left(\log\left(\frac{1}{1+e^{-\xi_{U_{il}}}}\right)-\log\left(\lambda_{U_{il}}\right)\right)+\left(1-\frac{1}{1+e^{-\xi_{U_{il}}}}\right)\times\left(\log\left(1-\frac{1}{1+e^{-\xi_{U_{il}}}}\right)-\log\left(1-\lambda_{U_{il}}\right)\right)\right] \\
&-\sum_{i=1}^{N}\sum_{l=1}^{L}\left[\left(\frac{1}{1+e^{-\xi_{U_{il}}}}\right)\times\left(\log\left(\beta^2_{0U_{il}}\right)-\log\left(\beta^2_{U_{il}}\right)+0.5\left(\alpha^2_{U_{il}}+\beta^2_{U_{il}}\right)/\beta^2_{0U_{il}}-0.5\right)\right] \\
&-\sum_{i=1}^{N}\left[\left(\frac{1}{1+e^{-\xi_{U_{i0}}}}\right)\times\left(\log\left(\frac{1}{1+e^{-\xi_{U_{i0}}}}\right)-\log\left(\lambda_{U_{i0}}\right)\right)+\left(1-\frac{1}{1+e^{-\xi_{U_{i0}}}}\right)\times\left(\log\left(1-\frac{1}{1+e^{-\xi_{U_{i0}}}}\right)-\log\left(1-\lambda_{U_{i0}}\right)\right)\right] \\
&-\sum_{i=1}^{N}\left[\left(\frac{1}{1+e^{-\xi_{U_{i0}}}}\right)\times\left(\log\left(\beta^2_{0U_{i0}}\right)-\log\left(\beta^2_{U_{i0}}\right)+0.5\left(\alpha^2_{U_{i0}}+\beta^2_{U_{i0}}\right)/\beta^2_{0U_{i0}}-0.5\right)\right]
\end{aligned} \tag{1}
$$

To compute the variational expectations $E_q[.]$ in equation (1), we use the properties of exponential family distribution. If variational distributions for $q\left(\mathbf{V}|\alpha_{\mathbf{V}},\beta^2_{\mathbf{V}}\right)$ and slab component $q\left(\mathbf{U}|\alpha_{\mathbf{U}},\beta^2_{\mathbf{U}}\right)$ are Gaussian distributions, then the exponential family representations are given by [1, 2]:

$$
\begin{aligned}
q\left(\mathbf{V}|\alpha_{\mathbf{V}},\beta^2_{\mathbf{V}}\right) &= 1/(\sqrt{2}\pi)\exp\left[(\alpha_{\mathbf{V}}/\beta^2_{\mathbf{V}})\mathbf{V} - 1/(2\beta^2_{\mathbf{V}})\mathbf{V}^2 - 1/(2\beta^2_{\mathbf{V}})\alpha^2_{\mathbf{V}} - \log(\beta_{\mathbf{V}})\right] \\
q\left(\mathbf{U}|\alpha_{\mathbf{U}},\beta^2_{\mathbf{U}}\right) &= 1/(\sqrt{2}\pi)\exp\left[(\alpha_{\mathbf{U}}/\beta^2_{\mathbf{U}})\mathbf{U} - 1/(2\beta^2_{\mathbf{U}})\mathbf{U}^2 - 1/(2\beta^2_{\mathbf{U}})\alpha^2_{\mathbf{U}} - \log(\beta_{\mathbf{U}})\right]
\end{aligned}
$$

So the natural parameters and sufficient statistics of the Gaussian distributions for $\mathbf{V}$ and $\mathbf{U}$ are $\eta_{\mathbf{V}} = \left[\alpha_{\mathbf{V}}/\beta^2_{\mathbf{V}}; -1/(2\beta^2_{\mathbf{V}})\right]$ , $T(\mathbf{V}) = \left[\mathbf{V};\mathbf{V}^2\right]$ and $\eta_{\mathbf{U}} = \left[\alpha_{\mathbf{U}}/\beta^2_{\mathbf{U}}; -1/(2\beta^2_{\mathbf{U}})\right]$ , $T(\mathbf{U}) = \left[\mathbf{U};\mathbf{U}^2\right]$, respectively.

Similarity, if variational distribution for $q\left(\boldsymbol{\gamma}|\boldsymbol{\xi}\right)$ is Bernoulli distribution, the exponential family representations are given by:

$$
q\left(\boldsymbol{\gamma}|\boldsymbol{\xi}\right) = \exp\left[\log\left(\boldsymbol{\xi}/(1-\boldsymbol{\xi})\right)\boldsymbol{\gamma} + \log\left(1-\boldsymbol{\xi}\right)\right]
$$

The natural parameters and sufficient statistics of the Bernoulli distributions for $\boldsymbol{\gamma}$ are $\eta_{\boldsymbol{\gamma}} = \left[\boldsymbol{\xi}/(1-\boldsymbol{\xi})\right]$ , $T(\boldsymbol{\gamma}) = [\boldsymbol{\gamma}]$.

#### 3.2 Reparameterization Trick

We use reparameterization trick to make the optimization of ELBO in equation (1). For the Gaussian distribution of the embedding matrix for metabolite abundances $\mathbf{V}$ that include weight matrix $V_{jl}$ and bias vector $V_{j0}$, we take samples via the following differentiable bi-variate transformation [3]:

$$
\begin{aligned}
V_{jl} &= \alpha_{V_{jl}} + \beta_{V_{jl}}\epsilon_{V_{jl}} \\
V_{j0} &= \alpha_{V_{j0}} + \beta_{V_{j0}}\epsilon_{V_{j0}}
\end{aligned}
$$

where $\epsilon_{V_{jl}} \sim \mathcal{N}(0,1)$ and $\epsilon_{V_{j0}} \sim \mathcal{N}(0,1)$. Then, we calculate and update their mean and standard deviation via the optimization of ELBO that can utilize the stochastic gradient approach as follows:

$$
\begin{aligned}
\nabla_{\alpha_{V_{jl}},\beta_{V_{jl}}}\mathcal{L}\left[\alpha_{V_{jl}},\beta_{V_{jl}}\right] &= \sum_{j=1}^{M}\sum_{l=1}^{L}\left(\nabla_{\alpha_{V_{jl}},\beta_{V_{jl}}}\mathrm{E_q}\left[\log\left(\mathrm{p}\left(\mathbf{D}\right)\right)\right] - \nabla_{\alpha_{\mathrm{V_{jl}}},\beta_{\mathrm{V_{jl}}}}\mathrm{KL}\left[\mathcal{N}\left(\alpha_{\mathrm{V_{jl}}},\beta^2_{\mathrm{V_{jl}}}\right)\|\mathcal{N}\left(0,\beta^2_{\mathrm{0V_{jl}}}\right)\right]\right) \\
\nabla_{\alpha_{V_{j0}},\beta_{V_{j0}}}\mathcal{L}\left[\alpha_{V_{j0}},\beta_{V_{j0}}\right] &= \sum_{j=1}^{M}\left(\nabla_{\alpha_{V_{j0}},\beta_{V_{j0}}}\mathrm{E_q}\left[\log\left(\mathrm{p}\left(\mathbf{D}\right)\right)\right] - \nabla_{\alpha_{\mathrm{V_{j0}}},\beta_{\mathrm{V_{j0}}}}\mathrm{KL}\left[\mathcal{N}\left(\alpha_{\mathrm{V_{j0}}},\beta^2_{\mathrm{V_{j0}}}\right)\|\mathcal{N}\left(0,\beta^2_{\mathrm{0V_{j0}}}\right)\right]\right)
\end{aligned}
$$

To reparameterize the discrete variable $\gamma$, we utilized the Gumbel-softmax approximation [4] to approximate Bernoulli distribution, we take samples of spike-and-slab distribution via the following transformation:

$$
\begin{aligned}
U_{il} &= \left(1+\exp\left(-\zeta_{\mathrm{U_{il}}}/\iota\right)\right)^{-1}\left(\alpha_{U_{il}} + \beta_{U_{il}}\epsilon_{U_{il}}\right) \\
\zeta_{U_{il}} &= \log\left(\xi_{\mathrm{il}}/\left(1-\xi_{\mathrm{il}}\right)\right) + \log\left(\kappa_{\mathrm{il}}/\left(1-\kappa_{\mathrm{il}}\right)\right) \\
U_{i0} &= \left(1+\exp\left(-\zeta_{\mathrm{U_{i0}}}/\iota\right)\right)^{-1}\left(\alpha_{U_{i0}} + \beta_{U_{i0}}\epsilon_{U_{i0}}\right) \\
\zeta_{U_{i0}} &= \log\left(\xi_{\mathrm{i0}}/\left(1-\xi_{\mathrm{i0}}\right)\right) + \log\left(\kappa_{\mathrm{i0}}/\left(1-\kappa_{\mathrm{i0}}\right)\right)
\end{aligned}
$$

where $\epsilon_{U_{jl}} \sim \mathcal{N}(0,1)$, $\epsilon_{U_{j0}} \sim \mathcal{N}(0,1)$, $\kappa_{il} \sim \mathcal{N}(0,1)$, and $\kappa_{i0} \sim \mathcal{N}(0,1)$. Then, we calculate and update the variational parameters via the optimization of ELBO that can utilize the stochastic gradient approach as follows:

$$\nabla_{\alpha_{U_{jl}},\beta_{U_{jl}},\zeta_{U_{il}}}\mathcal{L}\left[\alpha_{U_{jl}},\beta_{U_{jl}},\zeta_{U_{il}}\right] = \\ \sum_{i=1}^{N}\sum_{l=1}^{L}\left(\nabla_{\alpha_{U_{il}},\beta_{U_{il}},\zeta_{U_{il}}}\mathrm{E_q}\left[\log\left(\mathrm{p}\left(\mathbf{D}\right)\right)\right] - \nabla_{\zeta_{\mathrm{U_{il}}}}\mathrm{KL}\left[\mathrm{q}\left(\gamma\right)\|\mathrm{p}\left(\gamma\right)\right] - \nabla_{\alpha_{\mathrm{U_{jl}}},\beta_{\mathrm{U_{jl}}}}\mathrm{KL}\left[\mathcal{N}\left(\alpha_{\mathrm{U_{jl}}},\beta^2_{\mathrm{U_{jl}}}\right)\|\mathcal{N}\left(0,\beta^2_{0\mathrm{U_{jl}}}\right)\right]\right)$$

$$\nabla_{\alpha_{U_{j0}},\beta_{U_{j0}},\zeta_{U_{i0}}}\mathcal{L}\left[\alpha_{U_{j0}},\beta_{U_{j0}},\zeta_{U_{i0}}\right] = \\ \sum_{i=1}^{N}\left(\nabla_{\alpha_{U_{i0}},\beta_{U_{i0}},\zeta_{U_{i0}}}\mathrm{E_q}\left[\log\left(\mathrm{p}\left(\mathbf{D}\right)\right)\right] - \nabla_{\zeta_{\mathrm{U_{i0}}}}\mathrm{KL}\left[\mathrm{q}\left(\gamma\right)\|\mathrm{p}\left(\gamma\right)\right] - \nabla_{\alpha_{\mathrm{U_{j0}}},\beta_{\mathrm{U_{j0}}}}\mathrm{KL}\left[\mathcal{N}\left(\alpha_{\mathrm{U_{j0}}},\beta^2_{\mathrm{U_{j0}}}\right)\|\mathcal{N}\left(0,\beta^2_{0\mathrm{U_{j0}}}\right)\right]\right)$$
